# Tandem Requirement for Full Renal D_1_R and D_5_R Activity

**DOI:** 10.1101/736611

**Authors:** Selim Rozyyev, Annabelle P. Crusan, Andrew C. Tiu, Julie A. Jurgens, Justin Michael B. Quion, Laureano D. Asico, Robin A. Felder, Van Anthony M. Villar

## Abstract

The peripheral dopaminergic system promotes the maintenance of blood pressure homeostasis by engendering natriuresis, mainly through the renal D_1_R and D_5_R receptors. This effect is most apparent under conditions of moderate body sodium excess. Human and rodent renal proximal tubules express both receptors, which share common structural features and pharmacological profiles. Genetic ablation of either receptor in the kidney results in hypertension in mice. In this study, we demonstrated that in renal proximal tubules, these two receptors colocalized, co-immunoprecipitated, co-segregated in lipid rafts, and heterodimerized with one another, which was enhanced by treatment with the D_1_R/D_5_R agonist fenoldopam (1 μM, 30 min). Gene silencing via antisense oligonucleotides in renal proximal tubule cells abrogated cAMP production and sodium transport in response to fenoldopam. Our results highlight the cooperation and co-dependence of these two receptors through heterodimerization in renal proximal tubule cells.

## INTRODUCTION

Dopamine is a catecholamine that serves two important roles in neurobiology: as an intermediate in the synthesis of the neurotransmitters norepinephrine and epinephrine from tyrosine, and as a neurotransmitter by itself. In the mammalian brain, dopamine controls a variety of functions, including cognition, emotion, endocrine regulation, food intake, locomotor activity, memory, motivation, and positive reinforcement (1, 2). In addition to dopamine’s role in neurotransmission, dopamine is now recognized as an important regulator of fluid and electrolyte balance and blood pressure (1, 3, 4). Centrally, dopamine controls systemic blood pressure through the regulation of fluid and sodium intake via “appetite” centers in the brain (5) and by direct action on cardiovascular centers (1, 6). Peripherally, blood pressure regulation by dopamine is achieved by its direct actions on the blood vessels, heart, intestines, and kidney. Dopamine can also regulate water and electrolyte balance, indirectly through the regulation of the synthesis and release of hormones and humoral agents such as aldosterone, catecholamines, endothelin, prolactin, proopiomelanocortin, renin, and vasopressin, among others (1, 3, 4, 6–9). In the kidney, dopamine engenders natriuresis by increasing renal blood flow and glomerular filtration rate, and inhibiting renal tubular sodium reabsorption (1, 3, 4, 6–9). Under conditions of moderate sodium excess, dopamine generated by the renal proximal tubule, independent of renal nerves, and acting on the five dopamine receptors is responsible for about 50-60% of sodium excretion (4, 8, 9).

The dopamine receptors are classified into two subfamilies: D_1_-like dopamine receptors (D_1_R and D_5_R) and D_2_-like dopamine receptors (D_2_R, D_3_R and D_4_R) and are expressed in different segments of the nephron (1, 3, 4, 6–8). The D_1_R and D_5_R share a high degree of sequence and structural homology (intronless coding region, 80% identity in their transmembrane domains, and short 3^rd^ intracellular loop) and have similar ligand binding profiles and coupling patterns to cellular signals. They couple to stimulatory G proteins Gα_s_ and Gα_olf_ and target the same effector proteins such as the sodium-potassium pump (Na^+^/K^+^-ATPase) and sodium-hydrogen exchanger 3 (NHE3) (8, 9). There are, however, differences between these receptors. The D_5_R is constitutively active, has about 10x higher affinity for dopamine compared with D_1_R, and requires N-linked glycosylation for its plasma membrane localization (10). We have reported that the silencing of either D_1_R or D_5_R blunted the stimulation of cAMP production and inhibition of sodium transport caused by the D_1_R/D_5_R agonist fenoldopam, in isolated renal proximal tubule cells (RPTCs) and kidneys of C57Bl/6 mice (11, 12). In this study, we tested the hypothesis and provided proof that the D_1_-like dopamine receptors form a functional complex for their full activity in renal epithelial cells.

## MATERIALS AND METHODS

### Constructs

The human wild-type D_1_R and D_5_R were derived from in-house cDNA clones. pBiFC-YN155 and pBiFC-YC155 vectors (13) were kindly shared by Tom K. Kerppola. The YN vector contains the N-terminal portion of EYFP (residues 1-155) and the YC vector contains the C-terminal portion (residues 156-241) of EYFP, both fused to the C-terminus of the protein of interest. D_1_R cDNA was subcloned into the pBiFC-YN vector, while D_5_R was into the pBiFC-YC vector. Both constructs were transfected into hRPTCs. Transfection of D_1_R-EYFP alone was used as negative control. Alternatively, the D_1_R and D_5_R were subcloned into pCDNA3-mRFP and -mGFP, respectively. The fidelity of the cloned inserts was verified by DNA sequencing, and their expression was evaluated by RT-PCR and immunoblotting.

### Cell Culture

Immortalized human RPTCs (hRPTCs), obtained from normotensive Caucasian males (first characterized in 14, 15), were grown in DMEM-F12 medium (#11039047, Thermo Fisher Scientific, Waltham, MA), supplemented with 10% fetal bovine serum (#10082147, Thermo Fisher Scientific, Waltham, MA), selenium (5 ng/mL), insulin (5 μg/mL), transferrin (5 μg/mL), hydrocortisone (36 ng/mL), triiodothyronine (4 pg/mL), and epidermal growth factor (10 ng/mL). The cells were fed with fresh growth medium every three days. When visually confluent, the cells were subcultured for use in experiments and dissociated using trypsin (0.05%) and ethylenediaminetetraacetic acid (EDTA; 0.02%). For confocal microscopy studies, the cells were transiently co-transfected with pBiFC-DRD1-YN155 and pBiFC-DRD5-YC155 or pcDNA-DRD1-mRFP and pcDNA-DRD5-mGFP and then were grown on cover slips or in the polarized state using Transwells^™^ (#CLS3493, MilliporeSigma, Burlington, MA). Cells with low passage number (<20 passages) were used to avoid the confounding effects of cellular senescence. The cells tested negative for *Mycoplasma* infection.

### Immunohistochemistry and Immunofluorescence

Human kidneys were obtained as fresh surgical specimens from kidneys not used for transplantation or from patients who had unilateral nephrectomy due to renal carcinoma or trauma (approved by the Institutional Review Board, (IRB), of the University of Virginia Health Sciences Center). The intact and visually normal portion of kidneys obtained as discarded tissues from nephrectomies performed for cancer or trauma, was used for our studies. Immediately after excision, the tissues were immersion-fixed in HistoChoice^®^ (#VWRVH120, VWR, Radnor, PA) for at least 48 hours, dehydrated through graded alcohol and xylene baths, embedded in paraffin, and sectioned at 4 μm. After deparaffinization, rehydration and antigen retrieval, the endogenous peroxidase was blocked using 0.3% hydrogen peroxide in methanol for 30 mins. After the blocking solution was blotted off the tissue slice, each section was incubated overnight at 4°C in one of the following solutions containing antibodies diluted 1:50 in PBS (pH 7.4): (1) IgG affinity-purified rabbit antibody, (2) IgG affinity-purified D_1_R or D_5_R antibody preadsorbed against their respective immunizing peptides. Preadsorption was accomplished by incubating the antibody overnight at 4°C with a 10-fold M excess of the immunizing peptide. After 2 washes in PBS, the immunostaining was detected with avidin-biotin immunoperoxidase reaction (#PK-6200, Vectastain ABC kit, Vector Laboratories, Burlingame, CA), followed by visualization with 3,3’-diaminobenzidine (#D12384, MilliporeSigma, Burlington, MA). Tissue sections were lightly counterstained with hematoxylin to show the nuclei and placed under coverslips.

Alternatively, 4-μm thick sections of non-pathological human kidneys (#NBP1-78269, Imgenex of Novus Biologicals, Centennial, CO) were commercially obtained for immunofluorescence. The sections were antigen-retrieved using heat and pressure prior to double-staining for D_1_R and D_5_R, using rabbit anti-D_1_R labeled with CF^™^568 Fluor (#92235, Biotium, Fermont, CA) and rabbit anti-D_5_R antibody and donkey anti-rabbit secondary antibody tagged with Alexa Fluor^™^ 488 (#R37118, ThermoFisher Scientific Waltham, MA). The tissues were counterstained with wheat germ agglutinin (WGA) tagged with Alexa Fluor^™^ 647 (#A-31573, ThermoFisher Scientific, Waltham, MA) and DAPI (#H-1500, Vector Laboratories, Burlingame, CA) to visualize the plasma membrane and nuclei, respectively. Coverslip was mounted using a drop of Fluoro-Gel (#17985, EMS. Hatfield, PA). Confocal and differential interference contrast (DIC) images of the cells and kidney sections were obtained (along the XYZ axes for the cells and XY axis for the kidney) sequentially in separate channels to avoid bleed-through, using Carl Zeiss LSM 510 META with 63X/1.4 NA oil immersion objectives. The images were processed using Zen 2011 software (Carl Zeiss Microscopy GmbH, Jena, Germany).

### Antisense Experiments

The effect of 50 nM propyne/phosphorothioate-modified antisense against human *DRD5* (5’-136 CAGCATTTCGGGCTGGAC153 -3’) (GenBank Accession No. M67439) and human *DRD1* (5’- 277 GGTGTTCAGAGTCCTCAT 294n -3’) (GenBank Accession No. X55760) transcripts were compared with scrambled sequence controls (5’-GTCGCCCGAGCTTATGGA-3’ for *DRD5* and 5’- GGGTACTCTCT ATATCGG-3’ for *DRD1*). hRPTCs were seeded on a 24-well plate at a density of 5 × 10^4^ cells/well. After an 18-hr incubation, the cells were treated with the oligonucleotide mixed with DOTAP liposomal transfection agent (35 μg/ml) (#11811177001, Roche Diagnostics GmbH, Mannheim, Germany) for 3 to 6 hrs (16). After the treatment, the culture medium was aspirated and the cells were washed twice with PBS, pH 7.4 (#10010049, Thermo Fisher Scientific, Waltham, MA). After the second wash, the cells were incubated with normal culture medium at 37°C in 95% air and 5% CO2 for a total treatment time of 72 hrs.

### Lipid Raft Isolation & Sucrose Gradient Ultracentrifugation

hRPTCs were grown to confluence in 15-cm dishes, washed with PBS, pH 7.4, scraped, pelleted, and spun for 1 min at 4°C. After discarding the supernatant, SPE Buffer I from the FOCUS^™^ SubCell kit (# 786-260, G-Biosciences, St Louis, MO) was added to the pellet. The samples were vortexed and incubated on ice for 10 min. The cells were then homogenized and spun to pellet sequentially the nuclei and mitochondria. The supernatant was transferred to a new tube and spun at 100,000 × *g* for 60 min at 4°C in a SW50.1 swinging bucket rotor (Beckman CoulterTM, Palo Alto, CA) to pellet the membrane fraction. The plasma membrane-enriched pellet was subjected to detergent-free sucrose gradient ultracentrifugation (17, 18). Twelve fractions were collected and immunoblotted for D_1_R, D_5_R, and caveolin-1.

### Co-Immunoprecipitation

Cell lysates from hRPTCs were prepared using RIPA lysis buffer (50 mM Tris-HCl pH 7.4, 150 mM NaCl, 1% Triton X-100, 1% sodium deoxycholate, 0.1% SDS, #89901, Thermo Fisher Scientific, Waltham, MA) supplemented with protease inhibitors (1 mM PMSF, 5 μg/ml aprotinin and 5 μg/ml leupeptin, #78443, Thermo Fisher Scientific, Waltham, MA). Fifty ul of Dynabeads Protein A/Protein G (#10015D, Thermofisher Scientific, Waltham, MA) was washed with PBST and then admixed with 2 μg each of the immunoprecipitating antibody or normal rabbit IgG for 10 min on rotation. The beads were then washed 3x with PBST and then admixed with equal amounts of cell lysates (300 μg protein) for 10 min on rotation. The bound proteins were eluted by the addition of Laemmli buffer. The samples were immunoblotted and visualized for proteins of interest via chemiluminescence (SuperSignal West Dura Substrate, Pierce Biotechnology, Inc.) and autoradiography. The immunoprecipitates were run alongside with total cell lysates as positive control. GAPDH was used as loading control.

### Immunoblotting

The proteins were resolved by 10% sodium dodecyl sulfate (SDS) polyacrylamide gel electrophoresis and electrophoretically transferred onto nitrocellulose membranes. The transblot sheets were blocked with 5% normal donkey serum in TBST buffer Tris-buffered saline (10 mM Tris-HCl pH 7.5, 75 mM NaCl, #9997S, Cell Signaling Technology) and 0.1 % Tween 20] and incubated with diluted primary antibody for 1 hr at room temperature or overnight at 4°C. The transblots were washed with TBST buffer 3x and then incubated with diluted secondary peroxidase-conjugated affinity-purified antibody for 1 hr at room temperature. The immunoblots were copiously washed and developed with Enhanced Chemiluminescence (ECL Western Blotting Detection Kit; Amersham). The immunoblots were quantified using Quantiscan (19).

Antibodies against D_1_R (#GTX53978) and D_5_R (#GTX47806) were purchased from GeneTex (Irvine, CA USA). Specificity of these antibodies were tested in mock- and *DRD1* and DRD5-silenced cells. Cell lysates were mixed with Laemmli sample buffer (#1610737, BIO-RAD, Hercules, CA) boiled for 5 min, subjected to electrophoresis on 10% sodium dodecyl sulfate (SDS) polyacrylamide gel and transferred electrophoretically onto nitrocellulose membranes. Non-specific binding was blocked with 5% non-fat dry milk in TBST buffer. The membrane was then probed with the primary antibody (anti-D_1_R or anti-D_5_R, diluted 1:500) for one hr. After three washes, the membrane was incubated with peroxidase-labeled donkey anti-rabbit IgG (#NA9340-1ML, Ge Healthcare Life Sciences, Chicago, IL, USA) with 5% nonfat dry milk for one hr. In some studies, the antibodies were pre-adsorbed with their respective immunizing peptide (1:5 wt/wt incubated overnight at 4 °C). Specific bands were visualized using ECL.

### cAMP Measurement

The cells washed twice with PBS, pH 7.4, starved with serum-free medium for 2 hrs and incubated in 400 μls of PBS containing 1 μM 3-isobutyl-1-methyl xanthine (IBMX, #I5879, MilliporeSigma, Burlington, MA) at 37°C for 30 min in the presence or absence the D_1_R/D_5_R agonist fenoldopam (1 μM) (20). The reaction was terminated by aspirating the medium and washing the cells twice with PBS, pH 7.4 and freezing them at −80°C for 1 hr. The cells were then lysed with 0.1N HCl and cAMP was measured by radioimmunoassay (16).

### siRNA-mediated Gene Silencing

hRPTCs from normotensive subjects were seeded at 6-well tissue culture plates (Corning Inc.) and incubated at 37 °C in 95% air/5% CO2 for 24 hrs and at the time they were 50% confluent, the cells were treated with 10 pmol/L of D_1_R, D_5_R, mixed D_1_R and D_5_R-specific siRNA (#SI00015792, #SI00015869, Qiagen Science Inc., Germantown, MD) or non-silencing “mock” siRNA. After 48 hrs of the treatment, the cells were washed with PBS, starved with serum-free medium for 2 hrs, followed by incubation with 1μM of IBMX at 37 °C for 15 mins in the presence or absence of the D_1_R/D_5_R agonist fenoldopam (1 μM, #1659, Tocris Bioscience, Bristol, UK) or forskolin (5 μM, #1099, Tocris Bioscience, Bristol, UK). At the end of the incubation period, the media were removed and the cells rinsed twice with PBS. The cAMP responses were then measured using The DetectX^®^ Cyclic AMP (cAMP) Direct Immunoassay kit (#K019-H1, Arbor Assay Inc., Ann Arbor, MI).

### Cellular Sodium Transport Studies

hPRTCs were grown to confluence under polarized conditions on Corning HTS Transwell^™^ support inserted in 12-well plates and incubated at 37 °C incubator with humidified 5% CO_2_ in 95% air. The cells were serum-starved for 2 hrs and treated with fenoldopam (1 μM, 30 min), or vehicle (PBS, pH7.4 as control), at the basolateral area. After washing, the cells were incubated with the permeant sodium indicator sodium green tretraacetate (5 μM, #S6901, Invitrogen, Carlsbad, CA) in complete medium without phenol red. The cells were washed with PBS and the fluorescence signal for each Transwell^™^ was measured using the VICTOR3 V^™^ Multilabel Counter (excitation 485 nm and emission 535 nm). Ouabain (50 μM, 1 hr, #1076, Tocris Bioscience, Bristol, UK), added to the basolateral area, was used to inhibit Na^+^/K^+^-ATPase activity.

### Animal Care

Adult (8 wk-old) male C57Bl/6J mice were obtained from Jackson Laboratory (#000664, Bar Harbor, ME). All animal experiments were conducted in accordance with U.S. National Institutes of Health guidelines for the ethical treatment and handling of animals in research and approved by the George Washington University Institutional Animal Care and Use Committee (IACUC). The mice were housed in a temperature-controlled facility with a 12:12-hr light-dark cycle and fed with mouse chow and water *ad libitum* for at least 2 wks before any studies were performed. Mice were placed under anesthesia (50 mg/kg, pentobarbital sodium, #76478-501-50, Akorn Inc., Lake Forest, IL). The kidneys were initially perfused with heparinized saline prior to extraction for sucrose gradient ultracentrifugation.

### Statistical Analysis

Numerical data are expressed as mean ± standard error of the mean (SEM). Significant difference between two groups was determined by Student’s *t*-test, while significant differences among groups (>2) were determined by one-way factorial ANOVA and corresponding post-hoc test. A P value less than 0.05 was considered significant. Statistical analysis was performed using SSPS.

## RESULTS

### Renal D_1_R and D_5_R share the similar anatomical distribution in kidney

The D_1_R was found mainly at the luminal and basolateral membranes of proximal tubules and collecting ducts (**Figure 1A**) while the D5R was observed mainly at the luminal membrane of proximal tubules and collecting ducts (**Figure 1C**) in the human renal cortex. The D5R was also expressed at the medullary thick ascending limb of Henle (mTAL) (**Figure 1E**). Since the D_1_-like receptors share almost the same anatomical distribution at the proximal tubules, we next determined whether the receptors interact in hRPTCs and observed that these receptors co-immunoprecipitated at the basal state (mostly as dimers) and upon treatment (both as monomers and dimers) with the D_1_-like (D_1_R/D_5_R) receptor agonist fenoldopam in D_5_R-enriched HEK293 cells (Novus Biologicals Cat. #NBL1-10019) and hRPTCs (**Figures 1I, 1J**).

**Figure 1.**
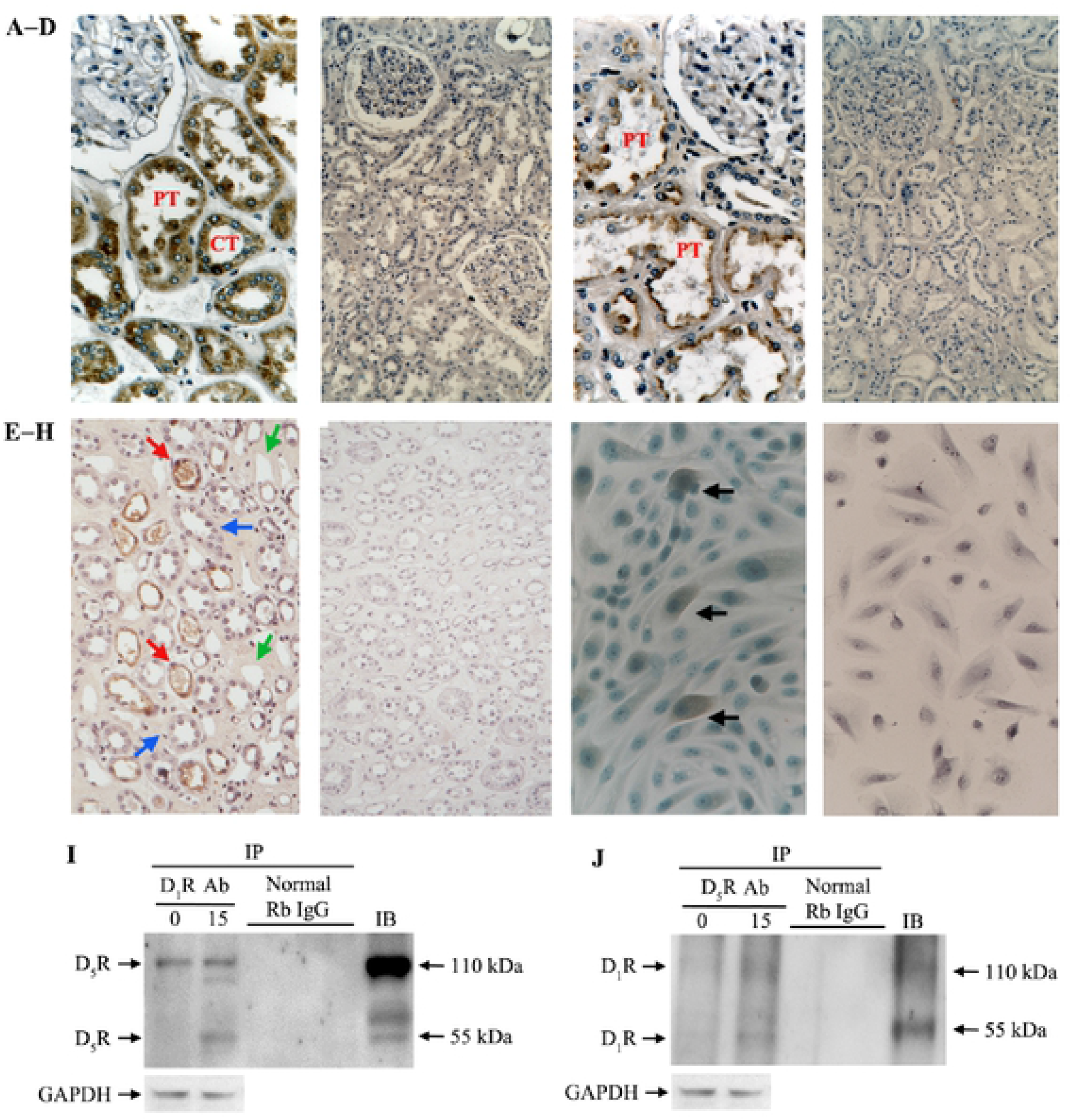
Expression and Interaction of Renal D_1_R and D_5_R. **A:** In the human renal cortex, D_1_R immunostaining was found throughout the cytoplasm, luminal, and basolateral membranes in renal proximal tubules (PT) and collecting tubule (CT). By contrast, **C:** D5R immunostaining was noted in luminal but not in basolateral membranes of renal proximal tubules (RPT) or collecting tubule (CT). **E:** D5R immunostaining was also noted in the medullary thick ascending limbs of Henle (red arrows) but not in medullary collecting ducts (blue arrows) or capillaries (green arrows). No staining was evident in slices with the D_1_R (**B**) and D_5_R (**D, F**) antibody preadsorbed with their respective immunizing peptides. D5R immunostaining was also noted in some of the cultured human renal proximal tubule cells (hRPTCs) (black arrows) (**G**). No staining was evident in cultured hRPTCs with the D_5_R antibody pre-adsorbed with the D_5_R immunizing peptide (**H**). Total cell lysates from hRPTCs treated with vehicle (0) or fenoldopam (1 μM/15 min) (18) were immunoprecipitated (IP) with anti-D_1_R or anti-D_5_R antibodies and immunoblotted for D_5_R or D_1_R, respectively (**I, J**). Normal rabbit (Rb) IgG was used as immunoprecipitant for the negative control. A D_5_R-enriched HEK-293 cell lysate (Novus Biologicals Cat. #NBL1- 10019) (I) or a hRPTC lysate was used for regular immunoblot (IB). (**J**)

We next evaluated the interaction between the receptors in renal epithelial cells and observed that the D_1_R-RFP and D_5_R-GFP were predominantly expressed at the plasma membrane of HEK-293 cells where they extensively colocalized at the basal state (**Figure 2A**). Specifically, both receptors were distributed at the apical and basolateral membranes where they strongly colocalized and partly at the cytoplasm of HEK-293 cells grown in the polarized state in Transwells^™^ (**Figure 2B**). We confirmed the physical interaction between these receptors by performing bimolecular fluorescence complementation (BiFC) and observed that the D_1_R and D_5_R basally interact in terminally differentiated hRPTCs, conceivably by forming heterodimers at the plasma membrane and cytoplasm (**Figure 2C**). Moreover, both receptors were robustly expressed at the proximal tubules where they predominantly colocalized at the brush borders in the human kidney (basally) and the mouse kidney (basally and upon fenoldopam stimulation) (**Figures 3A, B**).

Many G protein-coupled receptors (GPCRs) reside in dynamic signaling platforms in specific plasma membrane microdomains, in lipid and non-lipid rafts (18, 21). Therefore, we also evaluated the membrane distribution of the D_1_R and D_5_R via sucrose gradient centrifugation and found that the D_1_Rs partitioned to both lipid and non-lipid raft microdomains, while the D5Rs were located mostly at the non clipid raft microdomains at the basal state of hRPTCs (**Figure 4**). Agonist treatment with fenoldopam resulted in more D_1_R and D_5_R localization to the lipid rafts, where the adenylyl cyclases 5/6 are located (22), although a large portion of these receptors remained in non-lipid rafts. Thus, our findings demonstrate the shared localization of both D_1_R and D_5_R in the plasma membrane of renal epithelial cells.

**Figure 2.**
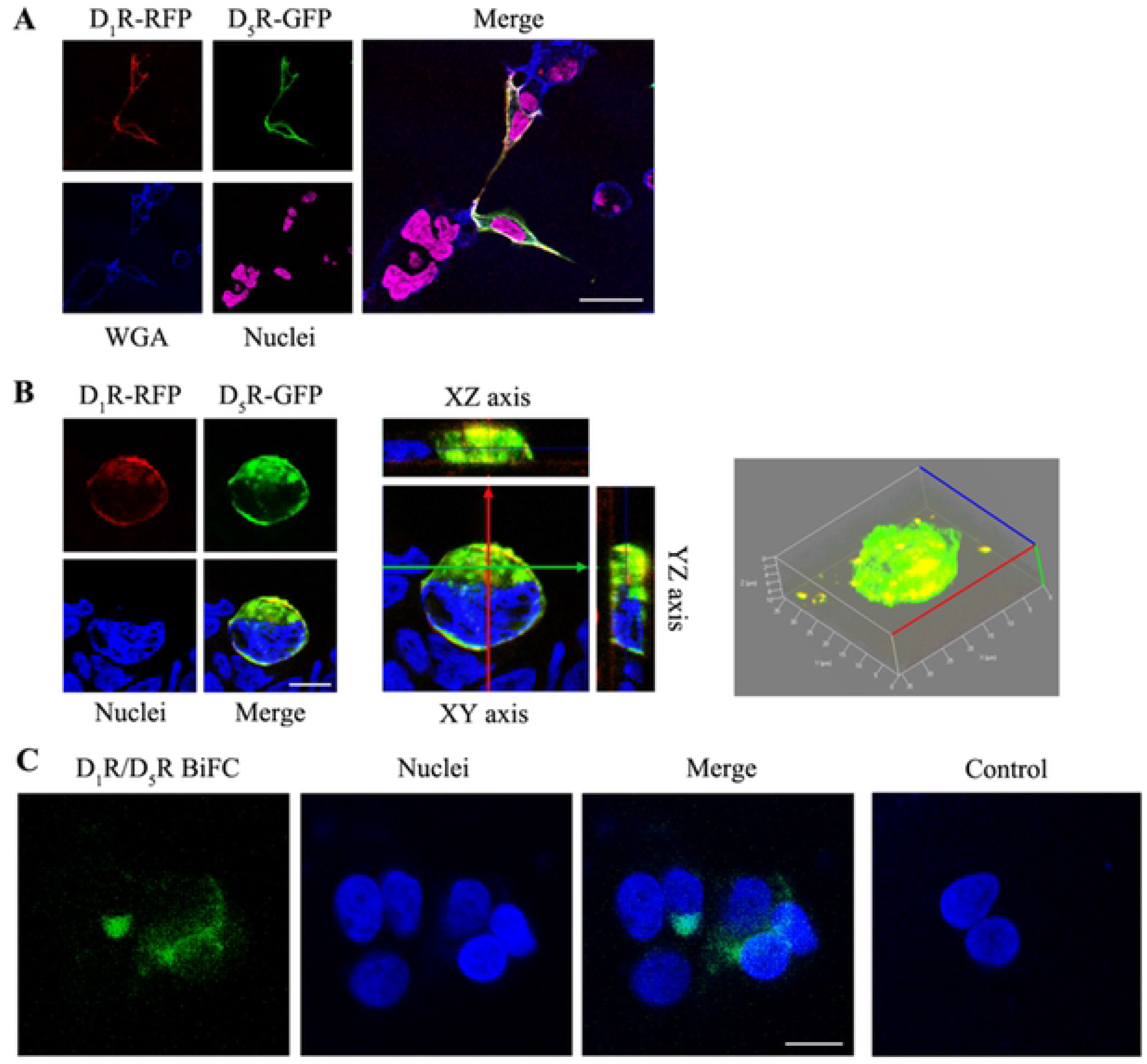
Colocalization of Renal D_1_R and D_5_R in Renal Epithelial Cells. **A & B**: HEK-293 cells were transiently transfected with D_1_R::RFP and D_5_R::GFP and then grown on a cover slip or in a polarized state in a Transwell^™^ insert; the colocalization of these receptors was evaluated via laser scanning confocal microscopy. Wheat germ agglutinin (WGA) tagged with Alexa Fluor□ 647 was used to visualized the plasma membrane, while DAPI (#H-1500, Vector Laboratories, Burlingame, CA) was used to visualized the nuclei. Colocalization was observed as discrete areas of white (red+green+blue) or yellow (red+green). Serial images along the X, Y, and Z axes were obtained in cells grown in Transwell^®^ and stitched together to show 3D colocalization of the receptors. 630x magnification, scale bar = 10 μm, n=3 independent experiments. **C**: hRPTCs were double-transfected with D_1_R and D_5_R tagged with the C and N termini of the fluorescent protein EYFP, respectively. The cells were grown on cover slips for 48 hours post-transfection, serum-starved for 2 hrs, and then prepared for confocal microscopy. The interaction of the tagged receptors results in the reconstitution and fluorescence of EYFP (pseudocolored green). An overlay of the BiFC signal and the nucleus (pseudocolored blue) is shown to indicate the distribution of the D_1_R-D_5_R complexes, 630x magnification, scale bar =10 μm, n=3 independent experiments. Transfection of D_1_R-EYFP alone was used as negative control.

**Figure 3.**
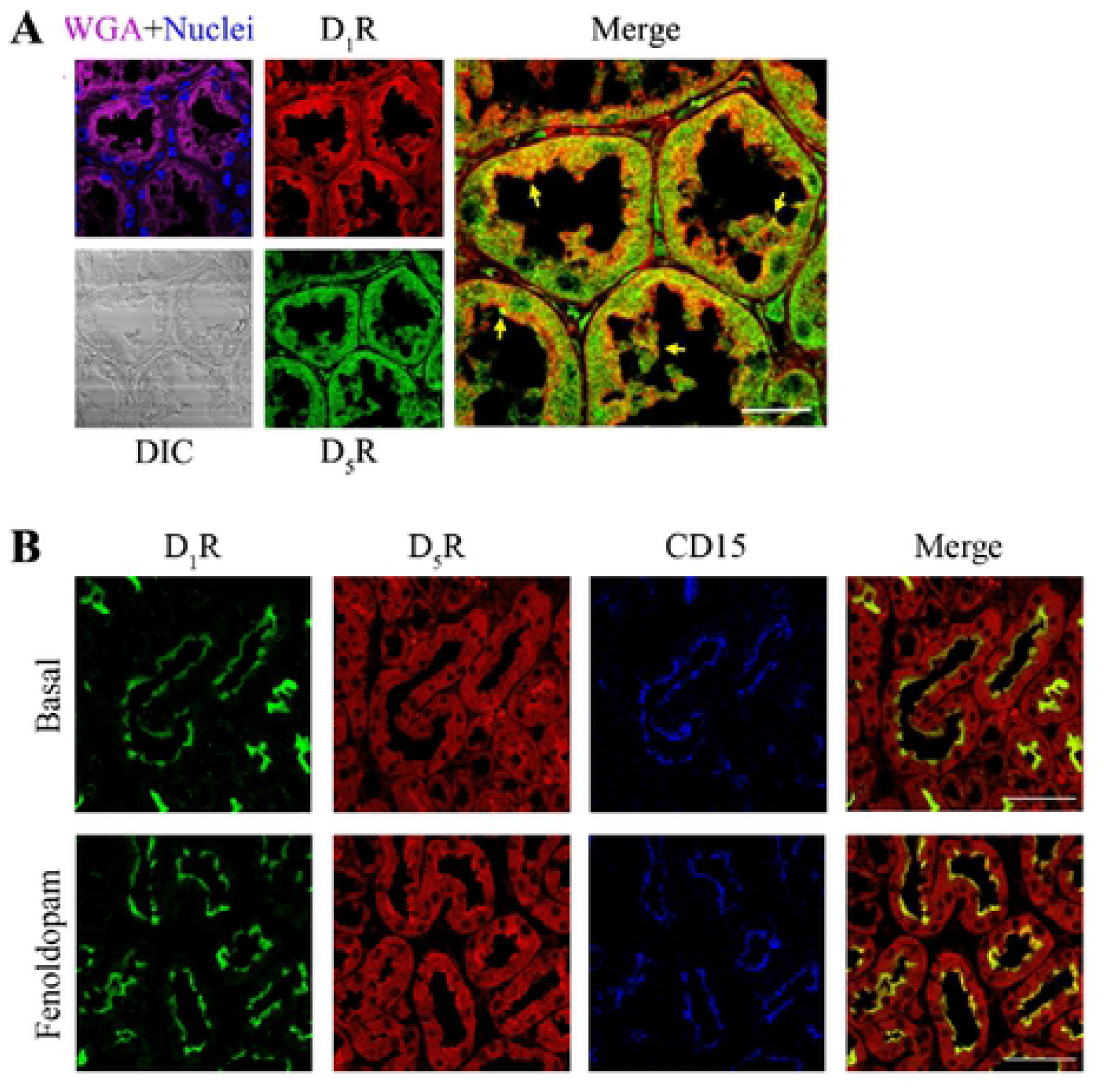
Colocalization of Renal D_1_R and D_5_R in the Kidney. A section of the human kidney was double immunostained for endogenous D_1_R (red) and D_5_R (green) (**A**). WGA (magenta) was used to visualize the plasma membrane, e.g., apical brush border of proximal tubules, while DAPI was used to visualize the nuclei. Colocalization in yellow is indicated by the arrows. DIC = differential interference microscopy. Sections of a mouse kidney infused with either vehicle (Basal) or fenoldopam was double immunostained for endogenous D_1_R (green) and D_5_R (red) (**B**). The RPT marker CD15 (23) and DAPI were used to visualize the brush border and nuclei, respectively. Colocalization is indicated by the yellow or white areas in merged images. 630x magnification, scale bar=10 μm, n=3 independent experiments.

**Figure 4.**
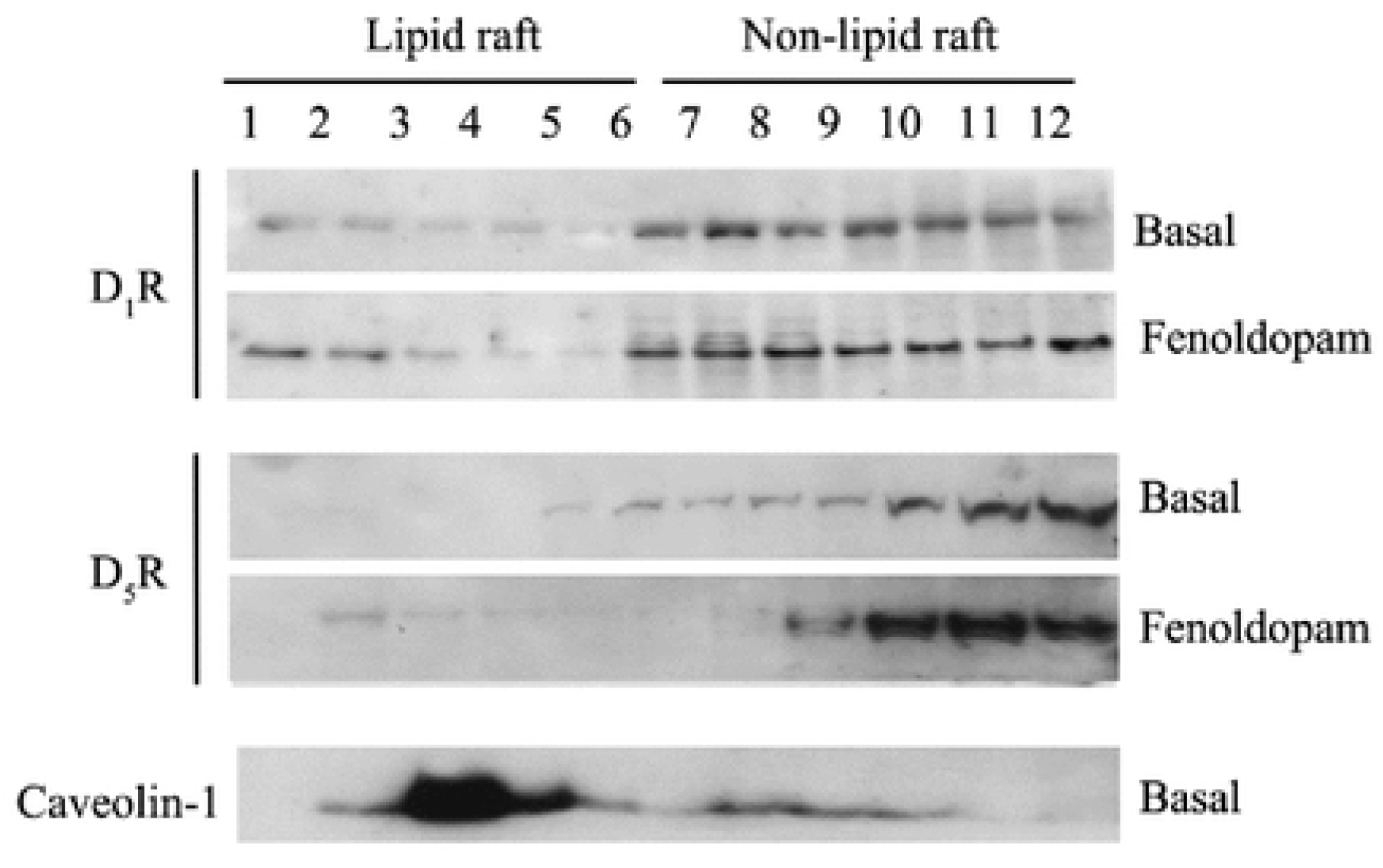
Co-segregation of Renal D_1_R and D_5_R in Lipid Rafts and Non-Lipid Rafts. The distribution of the D1-like dopamine receptors at the basal state and after fenoldopam treatment (1 μM, 15 min) in lipid rafts (more buoyant fractions 1-6) and nonlipid rafts (fractions 7-12) of hRPTCs was evaluated via detergent-free sucrose gradient ultra-centrifugation. Caveolin-1 is a lipid raft marker. n=3 independent experiments.

### D_1_R or D_5_R deficiency diminishes the activity of the other receptor

We next determined the functional relevance of the ability of the renal D_1_R and D_5_R to heterodimerize. Human renal proximal tubules cells (hRPTCs) were grown in Transwells^™^ and *DRD1, DRD5*, or both *DRD1* and *DRD5* genes were silenced via antisense RNA. hRPTCs were transfected with antisense or scrambled oligonucleotides or vehicle and then treated with either fenoldopam (1 μM, 30 min) or vehicle (basal) as negative control. The use of antisense oligonucleotides resulted in approximately a 25% (*DRD1*) or 50% (*DRD5*) reduction in the number of endogenous receptors expressed (Figure 5A). The cAMP response to fenoldopam treatment was blunted by ~30% (*DRD1*) or 70% (*DRD5*) when either receptor was depleted or by ~90% when both receptors were depleted (Figure 5B), indicating the requirement of both receptors for their full activity. Pre-treatment with either vehicle or scrambled oligonucleotides resulted in the expected increase in cAMP production upon fenoldopam treatment; the siRNA-mediated inhibition of the fenoldopam-mediated increase in cAMP production is greater with the silencing of both receptors than the silencing of either receptor alone (**Figure 5C**).

**Figure 5.**
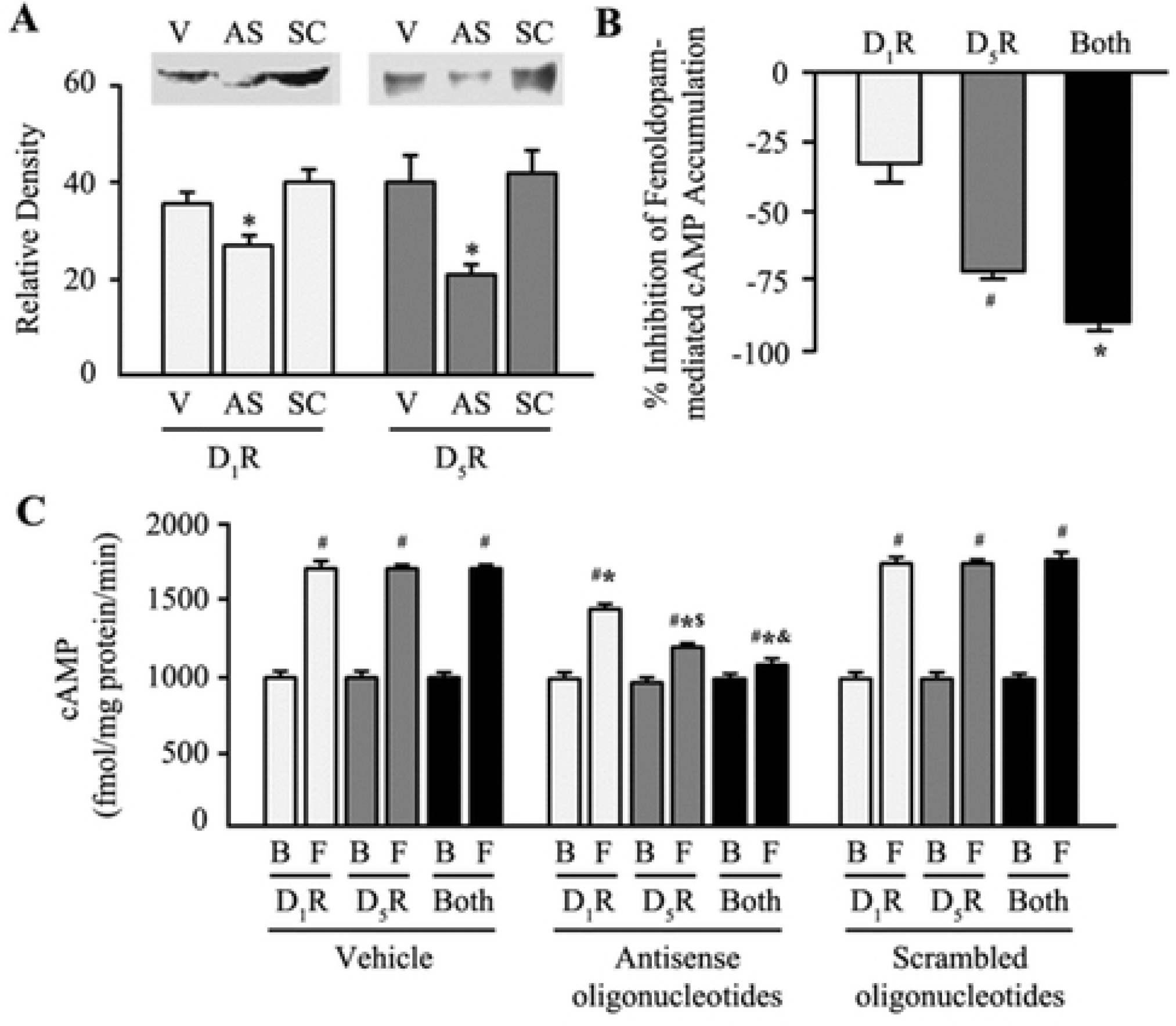
cAMP Production in Response to D_1_R/D_5_R Gene Silencing via Antisense Oligonucleotides. **A**: Effect of D_1_R or D_5_R antisense or scrambled oligonucleotides on D_1_R or D_5_R expression. *P<0.05 *vs.* vehicle or scrambled oligonucleotides, one-way ANOVA and Newman-Keuls post-hoc test. Immunoblots are shown in the inset. V = vehicle-treated cells, AS = Antisense oligonucleotide-treated cells, SC = scrambled oligonucleotide treated cells. **B**: Change in fenoldopam-stimulated cAMP accumulation in hRPTCs caused by antisense oligonucleotides against *DRD1* and *DRD5* mRNA. #P<0.05 *vs*. D_5_R, *P<0.05 vs. D_1_R or D_5_R, one-way ANOVA and Newman-Keuls post-hoc test, n = 3/group. **C**: Effect of D_1_R or D_5_R antisense or scrambled oligonucleotides on D1-like agonist fenoldopam (1 μM, 30 min)-stimulated cAMP accumulation in hRPTCs. #P<0.05 *vs*. Basal (B), *P<0.05 vs. fenoldopam (F) vehicle or scrambled, $P<0.05 vs. others Antisense oligonucleotides, <0.05 vs. others Antisense oligonucleotides, one-way ANOVA and Newman-Keuls post-hoc test, n=3/group.

**Figure 6.**
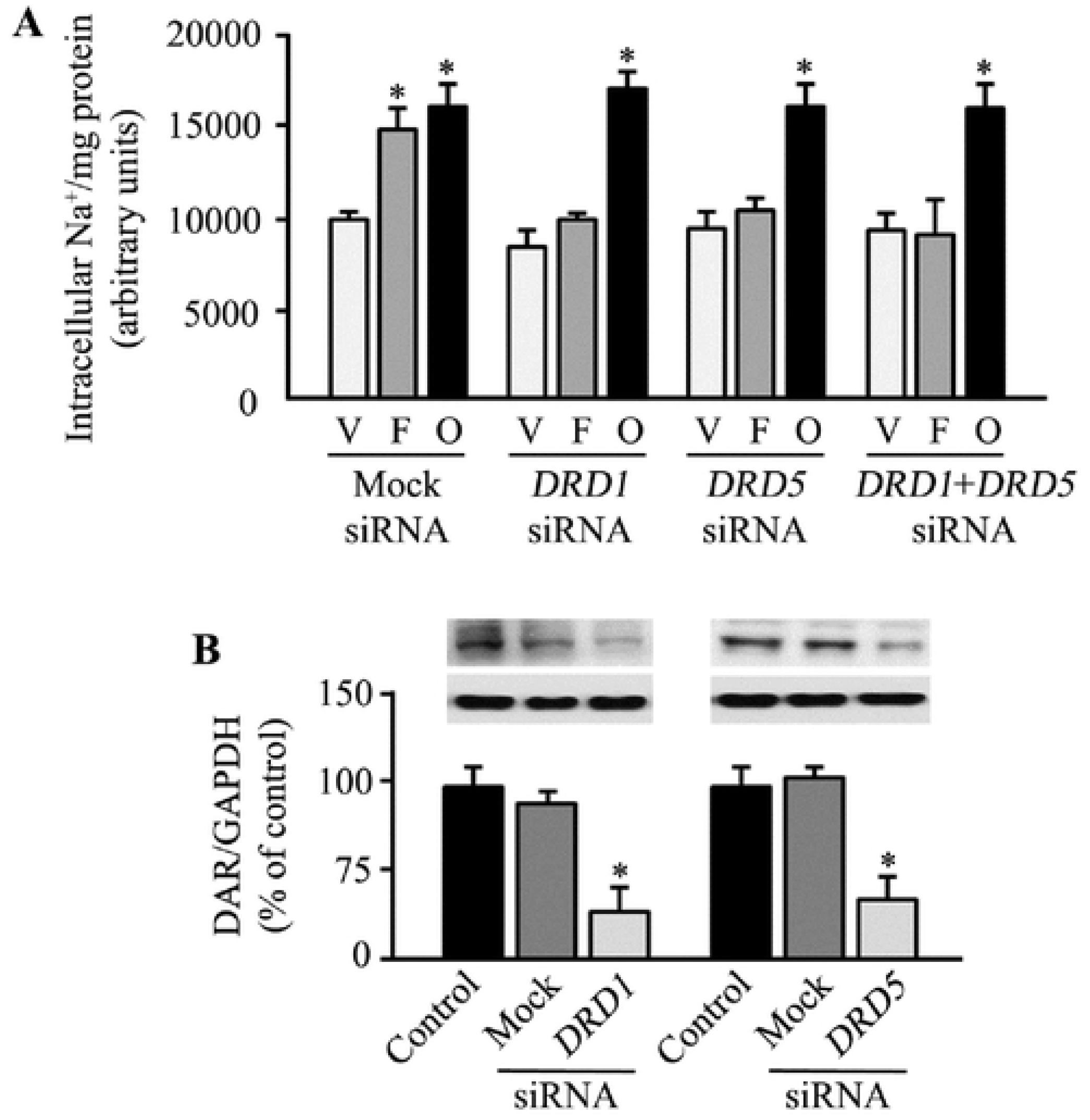
Sodium Transport in Response to D1R/D5R Silencing via siRNA. **A**: Intracellular Na^+^ accumulation in D_1_R- and/or D_5_R-depleted hRPTCs grown in the polarized state in Transwell^™^ inserts and treated with vehicle (V) or the D_1_R/D_5_R agonist fenoldopam (F; 1 μM, 30 min) or ouabain (O; 500 nM, 30 min) given at the basolateral compartment. *P<0.05, vs. others, one-way ANOVA and Holm-Sidak post-hoc test, n=3-4/group. **B**: The protein expression of D_1_R and D_5_R after 72 hr of receptor silencing through siRNA in hRPTCs. Non-silencing “Mock” siRNA was used as control. *P<0.05, vs. others, one-way ANOVA and Holm-Sidak post-test, n=3-4/group. GAPDH was used as housekeeping protein. Immunoblots are shown in the inset. DAR = dopamine receptor

We also evaluated the effect of receptor depletion through siRNA treatment on the ability of fenoldopam to inhibit sodium transport by the Na^+^/K^+^-ATPase. Typically, three Na^+^ ions are pumped out of the cell, in exchange for two K^+^ ion pumped into the cell across the basolateral membrane. D_1_R- or D_5_R-deficient hRPTCs were grown in the polarized state in Transwells^™^ and Na^+^/K^+^-ATPase activity was measured at the basolateral membrane. In mock siRNA-treated control cells, fenoldopam stimulation at the basolateral membrane side increased intracellular sodium, indicating the ability of activated receptors to inhibit Na^+^/K^+^-ATPase activity or sodium transport. In the cells with 90% depletion of D_1_R or 85% D_5_R depletion, fenoldopam was completely unable to increase intracellular sodium, indicating that the incomplete inhibition of the expression of either *DRD1* or *DRD5* nevertheless led to the complete failure to inhibit Na^+^/K^+^-ATPase activity. This more “complete” effect of fenoldopam on Na^+^/K^+^-ATPase activity as compared with cAMP production could be related to the greater decrease in D_1_R and D_5_R expression in the Na^+^/K^+^-ATPase activity relative to the cAMP study. These data indicate that the absence of one D_1_-like receptor completely prevents this action of the other D_1_-like receptor (**Figures 6A, B**). Pretreatment with the Na^+^/K^+^-ATPase inhibitor ouabain, as expected, increased intracellular sodium (**Figure 6A**), regardless of receptor expression level (**Figure 6B**), indicating that Na^+^/K^+^-ATPase is functioning normally, in the absence of the siRNAs against *DRD1* and/or *DRD5*.

## DISCUSSION

The homeostatic regulation of sodium balance and blood pressure hinges on the dynamic interaction among several cell surface receptors, usually belonging to different receptor families and subserving various, often antagonistic, effects on cellular processes. However, a number of receptors interact to promote, augment, or complete one another’s activity. Such is the case of the renal dopamine receptors on their ability to regulate tubular and vascular processes. Although the D_1_-like and D_2_-like dopamine receptors elicit opposite effects on cAMP production when individually stimulated, receptor occupation by their natural ligand dopamine results in the activation of Gα_s_/cAMP/ Protein Kinase A (PKA) pathway followed by the inhibition of sodium transport and natriuresis (1, 20, 24, 25). Moreover, genetic ablation of just one dopamine receptor subtype results in hypertension (1, 4, 12, 26, 27). The depletion or dysfunction of either the D_1_R or D_5_R also results in the abrogation of D_1_-like dopamine receptor activity even after agonist stimulation (1, 11, 12), suggesting a strict requirement for the expression of all dopamine receptor subtypes to be present and interact for their full renal activity. Additionally, there is evidence that the dopamine receptor subtypes negatively regulate the angiotensin II type 1 receptor and vice-versa (28–36). By contrast, dopamine receptor, e.g., D_1_R (37), D_2_R (38), and D_3_R (39) and angiotensin II type 2 receptor may positively regulate each other in the brain and kidney, effects in the latter organ have been shown to contribute to natriuresis and a further decrease in blood pressure. An aberrant dopamine receptor expression and/or activity may lead to increased blood pressure and hypertension (1, 3, 4, 6–9, 12, 27, 31, 34, 40–43).

The concept of protein clustering to form higher order complexes has been fairly established to explain the functional crosstalk among receptors, including the GPCRs (44). This organization profoundly influences protein function and many GPCR heteromers have been shown to assume distinct functional properties compared with the corresponding monomers. The ability of these receptors, including the renal D1R and D5R, to organize into functional oligomeric complexes requires several crucial elements to be present, including (a) common anatomical distribution, (b) physical interaction, direct or otherwise through adaptor or scaffold proteins, (c) shared signaling pathways, and (d) common biological triggers (e.g., high salt diet, dopamine) and endpoints (e.g., sodium transporters). We now demonstrate the presence of these crucial prerequisites for the cooperative activity of D_1_R and D_5_R. The D_1_-like receptors are expressed in not just in the same segments of the nephron, but these also partition to the same membrane microdomains, i.e., lipid rafts, upon agonist stimulation where they conceivably form functional signaling complexes. Moreover, these receptors physically interact basally and more so upon agonist stimulation, and are functionally co-dependent on one another to inhibit sodium transport (20, 37). Our data demonstrate the requirement for concurrent activation of both D_1_R and D_5_R, leading to heterodimerization and inhibition of RPT sodium transport, and by inference, blood pressure regulation.

The D_1_R is expressed in luminal, basolateral, and intracellular areas of the RPT and collecting ducts of mouse, rat, and human kidneys (1, 3, 4, 6–9, 19, 22, 24, 27–29, 31, 34, 36, 37, 41, 43, 45), whereas the D_5_R is seen only in the luminal membrane of RPTs and collecting ducts (1, 12, 26). The D_5_R but not the D1R receptor is visualized in medullary thick ascending limb of Henle (mTAL) of the human kidney, similar to studies in rat kidneys (1, 26).

Both the D_1_R and D_5_R are expressed in cultured hRPTCs (12, 20, 22, 24, 37, 43), similar to the porcine RPTCs (LLCPK1) (46), both of which are fully differentiated renal epithelial cells. Both receptors are functional in hRPTCs, and share some common signaling cascades. Both the D_1_R and D_5_R are linked to Gα_s_ and the stimulation of adenylyl cyclase activity (1, 3, 4, 6–9, 20, 22, 25). In renal cortical membranes, these receptors are also linked to Gα_q_ (1, 3, 4, 6–9, 20). We have reported that in hRPTCs, most of the stimulatory effect of fenoldopam on cAMP production can be accounted for by D_1_R; the *DRD1*-siRNA-mediated decrease in D_1_R expression was associated with a 90% inhibition of fenoldopam-mediated (1 μM/20 min) increase in cAMP accumulation (20). By contrast, a 75% decrease in D_5_R expression minimally affected fenoldopam-mediated stimulation of cAMP accumulation, indicating that the D5R under in hRPTCs these circumstances minimally affects cAMP production (20). Our current study in which 50% of D_5_R was silenced, there was a 75% decrease in fenoldopam-mediated stimulation of cAMP suggesting that the D5R can stimulate cAMP production. The difference between the previous study (20) and our current study could be related to the duration of incubation of fenoldopam (20 vs 30 min); we have reported that in hRPTCs, 1 μM fenoldopam increases cAMP production by about 53% from 20 to 30 min of incubation. Whether or not this can explain the difference in cAMP production between the current and previous study (20) remains to be determined. In the same study (20) in hRPTCs, the D5R couples with the Gαq and stimulates phospholipase C (PLC) and protein kinase C (PKC), and that both cAMP and PLC pathways inhibit Na^+^/K^+^-ATPase activity (20). The current study supports the previous study of Gildea et al (20); genetic silencing of either D_1_R or D_5_R impairs the ability of fenoldopam to inhibit Na^+^/K^+^-ATPase activity in hRPTCs.

The ability of GPCRs, in general, and of the dopamine receptors, in particular, to oligomerize is well established (47–56). These complexes may be composed of identical protomers or monomers (aka homomers) or of different protomers (aka heteromers). Oligodimerization facilitates proper ligand binding, efficient receptor endocytosis, and effective signal transduction, as well as modifies the desensitization profile and post-endocytic fate of the receptors (49). Between the D_1_-like and D_2_-like dopamine receptors, the D2-like dopamine receptors have been more frequently reported to oligomerize, either with one another (48, 50–52) or with other GPCRs, including the adenosine A1 and A2 receptors, ghrelin receptor, and the somatostatin receptor 2, among others (48, 53, 54). These oligomers have been described mostly in the central nervous system. The D_1_R couples with the D_3_R and results in synergism (47, 55), although these dopamine receptors have contrasting effects on cAMP when individually activated (1, 3, 4, 6–9). The D_1_R also heterodimerizes with adenosine A1 receptor (56) and the nonopioid σ-1 receptor (57) in the brain. By contrast, the D_5_R has not been described to oligomerize with other receptors, even in dopaminergic neurons. We now report that the D_1_R and D_5_R form functional oligomers in the renal parenchyma, specifically in renal proximal tubules, and that the absence of one blunts the effect of the other. Our observations do not only offer a mechanistic backdrop for the co-dependence between the two D_1_-like dopamine receptors, but also offer insights into the differences in the pharmacological profiles of antihypertensive agents at the cellular level. It is conceivable that the D_1_R-D_5_R heteromers are structurally and functionally expedient considering the shared ligand binding, signaling cascades, and biological targets for their actions.

Taken together, the current study underscores the requirement for agonist-activated D_1_R and D_5_R in lipid raft membrane microdomains to heterodimerize to achieve their full biological activity, i.e., control of sodium and fluid balance and vasomotor tone, which are two cornerstones in blood pressure regulation (**Figure 7**). The absence of one receptor affects the function of the other receptor and has dire functional consequences in the kidney, among others, and results in blunted natriuresis and elevation of blood pressure.

**Figure 7.**
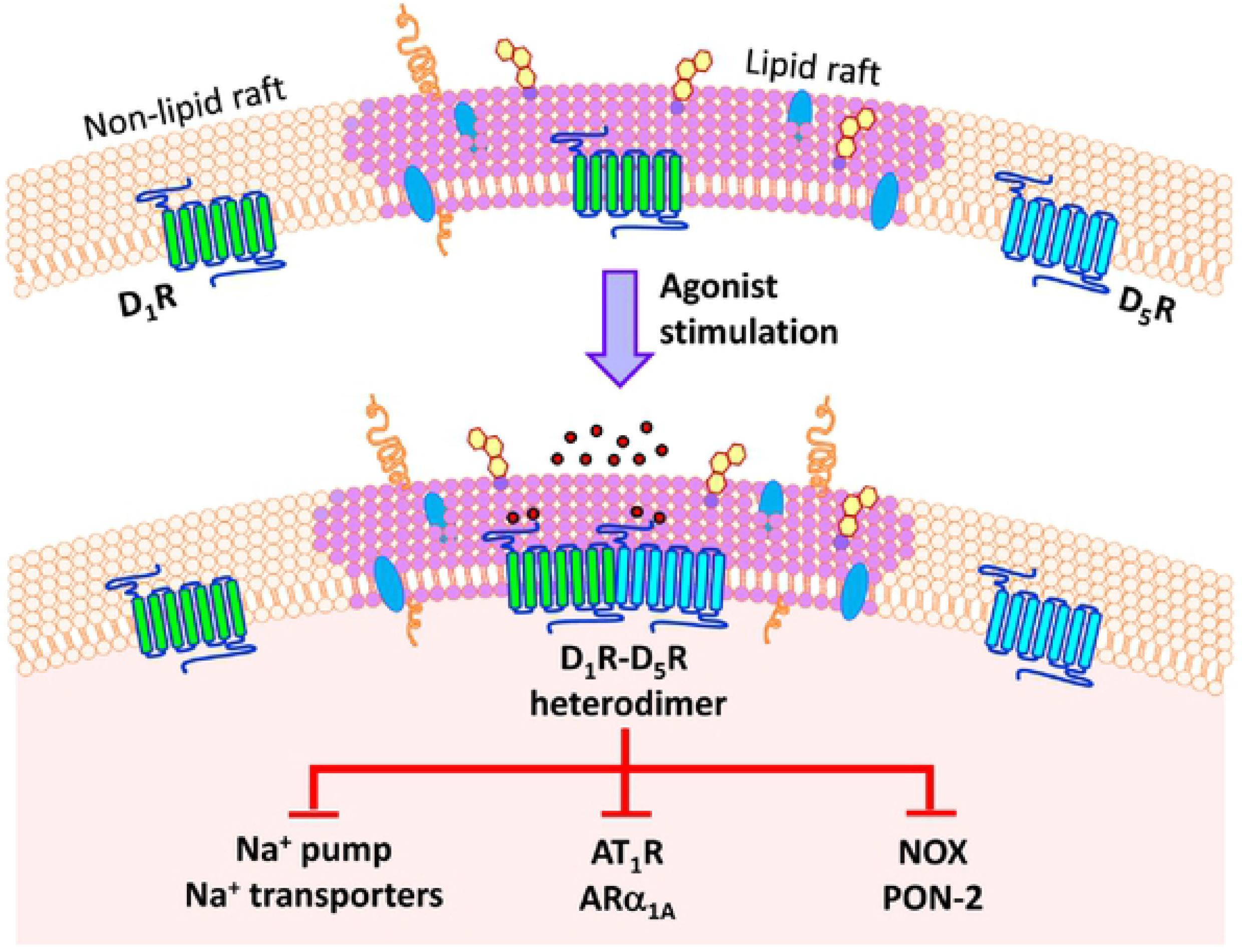
D_1_R and D_5_R work in concert in the kidney. Basally, the D_1_R is found in both lipid and non-lipid rafts, while the D_5_R is mostly in the non-lipid rafts. Agonist stimulation of both receptors result in the enrichment of both receptors in lipid raft microdomains where they conceivably form functional heterodimers to inhibit the sodium pump and sodium transporters (22, 24, 58), the AT_1_R (29, 34, 35, 41, 43, 59), ARα1A (60), and NOX but stimulate PON2 (61, 62), among others, to promote natriuresis and prevent the unfettered increase in blood pressure.

## ACKNOWLEDGMENTS

The pBiFC-YN155 and pBiFC-YC155 vectors were kindly shared by Professor Tom K. Kerppola of the University of Michigan Medical School.

## FUNDING SOURCES

The studies were supported, in part, by grants from the National Institutes of Health (5R01DK039308-32 and 5P01HL074940-13) and minigrants from the National Kidney Foundation of Maryland (NKF-MD) for VA Villar and LD Asico.

